# Small heat shock protein HSPB5 uses disorder to bind zinc with high affinity

**DOI:** 10.1101/2025.10.17.683119

**Authors:** Maria K. Janowska, Vanesa Racigh, Christoper N. Woods, María Silvina Fornasari, Rachel E. Klevit

## Abstract

Zinc is an essential metal that supports diverse cellular functions. Zinc exerts its biological activity through protein binding, serving as catalytic cofactors and structural stabilizers of many enzymes, transcription factors, and ubiquitin E3 ligases, among others. Despite total cellular zinc concentrations reaching hundreds of micromolar, free zinc levels are tightly buffered. Elevated free zinc promotes mismetalation and protein aggregation. While zinc is redox-inert, its cysteine-based protein ligands are readily oxidized. Oxidative modification of cysteines leads to zinc dissociation and a rapid increase in free zinc. With ∼3000 proteins in the human zinc proteome, uncontrolled zinc release could be highly deleterious. Metallothioneins buffer zinc under basal conditions, but their re-synthesis following oxidative inactivation occurs on the scale of hours, raising the question of how free zinc is managed in the interim. Histidine, the second most prevalent zinc-coordinating residue, is resistant to oxidative modification. We characterized zinc binding by the small heat shock protein HSPB5 (αB-crystallin), a cysteine-free, histidine-rich protein chaperone that responds to cellular stress and found: (1) HSPB5 binds zinc with high affinity and rapid reversibility; (2) zinc binding requires the disordered HSPB5 N-terminal region; (3) zinc binding increases HSPB5 disorder; and (4) prolonged zinc exposure promotes formation of assemblies of oligomers cross-bridged by zinc. We propose that HSPB5 has evolved specialized zinc-dependent properties distinct among human sHSPs, enabling it to function not only as a protein chaperone but also as a conditional zinc reservoir under oxidative stress.

## Introduction

Zinc is one of ten essential metals in humans, supporting processes such as immunity, oxidative stress responses, DNA repair, and aging (1–4). Unlike redox-active metals, zinc is chemically inert and its functions arise from protein binding. About 3000 human proteins—roughly 10% of the proteome— require zinc as a cofactor or structural stabilizer (5, 6). Although total intracellular zinc reaches hundreds of micromolar, free zinc is maintained at pM–nM levels mainly by metallothioneins, cysteine-rich proteins that bind zinc with high affinity (5, 7, 8). Under oxidative stress, cysteine thiolates can be oxidized, releasing zinc and elevating free zinc to levels that can drive mismetalation and aggregation (9–12). Such dyshomeostasis is implicated in disease, including Alzheimer’s plaques and cataractous lenses (13, 14). In proteins, zinc typically coordinates cysteine (Cys) and histidine (His) ligands (5, 6, 15, 16). Because Cys is redox-sensitive but His is not, His-rich, Cys-free proteins may serve as conditional zinc buffers when metallothioneins fail.

The small heat shock protein (sHSP) HSPB5 (αB-crystallin) is a compelling candidate for such a role. It is abundant, constitutively and ubiquitously expressed, and acts as a molecular chaperone that protects proteins from aggregation under stress (17–21). Notably, HSPB5 is enriched in His content with 9 His out of 175 residues (5.1%), compared with a 2.6% proteome-wide average (22). Like other sHSPs, it consists of an intrinsically disordered N-terminal region (NTR), a structured α-crystallin domain (ACD), and a disordered C-terminal region (CTR) (23–25). Interactions among structured ACD dimers, NTRs, and CTRs drive formation of polydisperse oligomers with ∼12-40 subunits, in which the intrinsically disordered regions exist in dynamic equilibrium between solvent-exposed and quasi-ordered states, a property central to sHSP function (21). A typical HSPB5 24-mer therefore contains over 200 His, making HSPB5 a plausible zinc-binding reservoir.

While zinc effects on HSPB5 have been reported (26–32), direct measurements of binding affinity, stoichiometry, and mechanistic consequences are lacking. We used biochemical and biophysical approaches to define zinc binding to HSPB5. We show that HSPB5 binds zinc with high (nanomolar) affinity and this requires the disordered NTR. Zinc induces at least two transitions, increases disorder within oligomers, and slows subunit exchange. Prolonged zinc exposure promotes formation of large species consistent with oligomers cross-bridged by zinc. The findings suggest that HSPB5 possesses specialized zinc-dependent properties unique among human sHSPs. We propose that, beyond its established role as a protein chaperone, HSPB5 may function as a conditional zinc reservoir during oxidative stress/zinc distress (33), when Cys-based sites release zinc. This dual role could link proteostasis with zinc buffering and highlights a previously unrecognized dimension of sHSP biology.

## Results

### HSPB5 binds Zn^2+^ with high affinity

To measure zinc binding, we used a competition assay between a molecule whose fluorescence is zinc-dependent and HSPB5 (34). FZ3 binds Zn^2+^ with a K_d_ of 9.1 nM (35) and fluoresces at ∼520 nm when bound to Zn^2+^. Under conditions of limiting zinc, addition of HSPB5 led to a loss of FZ3 fluorescence, indicating HSPB5 competes for zinc (Fig. 1A&B, 1^st^ column). Fitting *F*(520nm) as a function of [HSPB5] to a simple binding competition equation yielded an apparent K_d_ of 19 nM. In contrast, no competition for zinc was detected for the ACD dimer, despite containing 5 His per subunit, (Fig. 1A&B, 2^nd^ column). HSPB4, a closely related sHSP that also forms large oligomers, contains 6 structurally analogous His to those in HSPB5 and a His in its CTR (Fig. S1). Despite its 7 His residues per subunit, HSPB4 did not compete with FZ3 for zinc (Fig. 1A&B, 3^rd^ column). Notably, high-affinity zinc binding is recapitulated in an HSPB4 construct to which 3 NTR His were introduced at positions analogous to HSPB5 positions (HSPB4-3His, Fig. 1A&B, 4^th^ column). In sum, the results reveal that oligomeric HSPB5 can out-compete FZ3 for zinc and this high-affinity binding requires the NTR.

**Figure 1.**
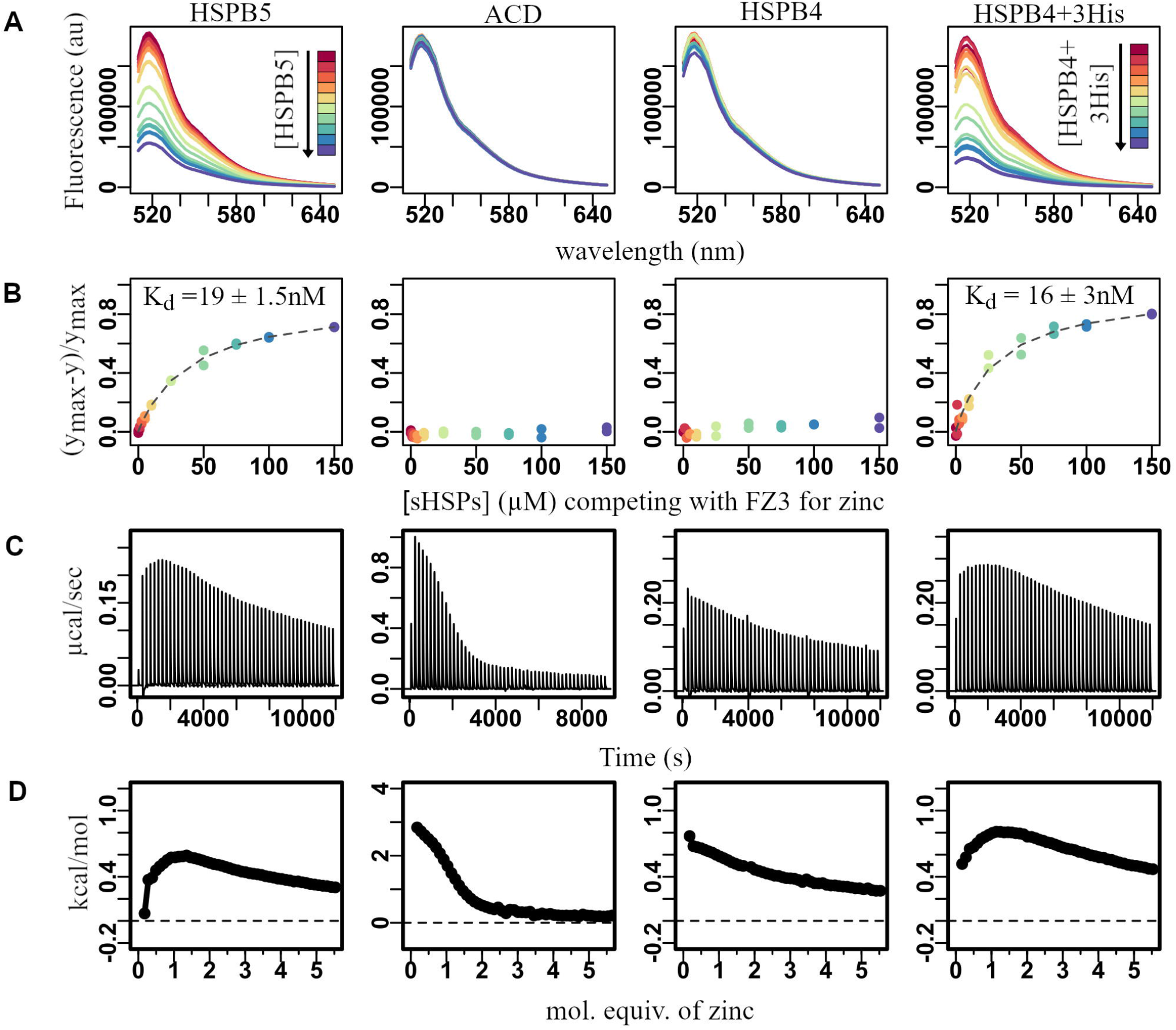
Zinc binding to HSPB5 depends on disordered NTR histidines. All panels (left to right) show data for HSPB5, HSPB5 ACD, HSPB4, and HSPB4 with introduction of 3 NTR His. A. Zinc competition assay. Fluorescence emission spectrum of FluoZin-3 (FZ3; 20 μM) in the presence of 1.5 μM zinc with increasing concentrations of sHSPs (0 → 150 μM). Colors correspond to increasing [HSPB5] from red to blue (rainbow color scheme). B. Fits of assays shown in part A. C. ITC injection profiles. D. ITC isotherms for addition of zinc to HSPB5 and HSPB4 constructs.

### HSPB5 undergoes multiple zinc-dependent transitions that depend on the disordered NTR

We used isothermal titration calorimetry (ITC) to assess zinc binding by HSPB5. Titrations were performed at 50 μM HSPB5, >10^3^-fold above the K_d_ measured by FZ3, such that stoichiometric binding should yield a linear, monophasic isotherm until saturation. Instead, the observed isotherm was biphasic: an exothermic phase at low zinc ratios (0–1 Zn^2+^/HSPB5) followed by an endothermic phase that saturated at >5 Zn^2+^/HSPB5 (Fig. S2, Table S1). In contrast, the HSPB5 ACD dimer produced a simple endothermic isotherm, confirming zinc binding but at much lower affinity than full-length oligomeric HSPB5. These observations suggest that at least two zinc-dependent processes occur in HSPB5 and implicate the disordered regions in driving the behavior.

Fitting the biphasic isotherm to a two-phase model revealed that both transitions are spontaneous at 25 °C (ΔG < 0). The first transition involved large decreases in both enthalpy (ΔH = –173 kJ/mol) and entropy (TΔS = –146 kJ/mol), while the second showed a small enthalpic gain (ΔH = 6 kJ/mol) and a modest entropic penalty (TΔS = –20.5 kJ/mol). Although these values do not permit detailed microscopic interpretation, they establish that zinc binding induces two thermodynamically distinct processes in HSPB5. For comparison, the ACD dimer fit to a single endothermic process (ΔH = 18 kJ/mol; TΔS = –47 kJ/mol), and HSPB4 oligomers also gave a monophasic, endothermic isotherm, confirming zinc binding with lower affinity than HSPB5 (Fig. 1C&D, Table S1).

To test the role of the NTR His residues, a single His was introduced into a Cys-free version of HSPB4 at positions analogous to HSPB5. Remarkably, introduction of His at either position 6 (Q6H) or 31 (G30H) produced HSPB4 variants that display biphasic isotherms (Fig. S3). The effect is position-specific, as substitution at position 18 (Y18H) retained the monophasic behavior of wild-type HSPB4. These results indicate that the presence of His at specific NTR positions is required for the biphasic zinc-dependent transitions observed in HSPB5. To probe the contribution of individual residues in HSPB5, each histidine was replaced with alanine and isotherms were collected for the resulting oligomers. All mutants retained biphasic behavior, showing that no single His is essential for this property (Fig. S4, S5).

Ultimately, ITC did not provide a clear stoichiometry of zinc binding to HSPB5 but instead revealed unusual zinc-dependent properties. First, although HSPB5 binds zinc with high affinity, it lacks discrete binding sites composed of defined residue sets. Second, the structured ACD dimer, with 10 His, binds zinc with substantially lower affinity than oligomeric HSPB5. Third, ITC detected two spontaneous zinc-dependent transitions. Because the observed heats cannot be directly ascribed to zinc binding under these conditions, we propose they reflect conformational reorganization within or among HSPB5 oligomers that involves the disordered NTR.

### Zinc-induced effects are dose-, time-, and temperature-dependent and fully reversible

HSPB5 oligomers increase in hydrodynamic radius in the presence of zinc (26). At 37 °C, dynamic light scattering (DLS) measurements showed an average radius of ∼7 nm under low zinc conditions, which remained stable without added zinc but increased over time to ∼8, 10, and 30 nm with 1×, 2×, and 10× zinc, respectively (Fig. 2A). Time courses fit to a first-order rate equation yielded condition-dependent rate constants and maximum sizes, with half-times of ∼60 and ∼90 min for 2× and 10× zinc (Fig. 2B). These effects occurred at 37 °C, but not at 25 °C (Fig. S6). Analytical size-exclusion chromatography confirmed these findings: (1) no elution change with 1× or 2× zinc at 25 °C; (2) zinc-dependent shifts after incubation at 37 °C; and (3) progressive shifts with longer incubation (Fig. S7A–C). Together, these results demonstrate that zinc-dependent changes in HSPB5 oligomer size are dose-, time-, and temperature-dependent.

**Figure 2.**
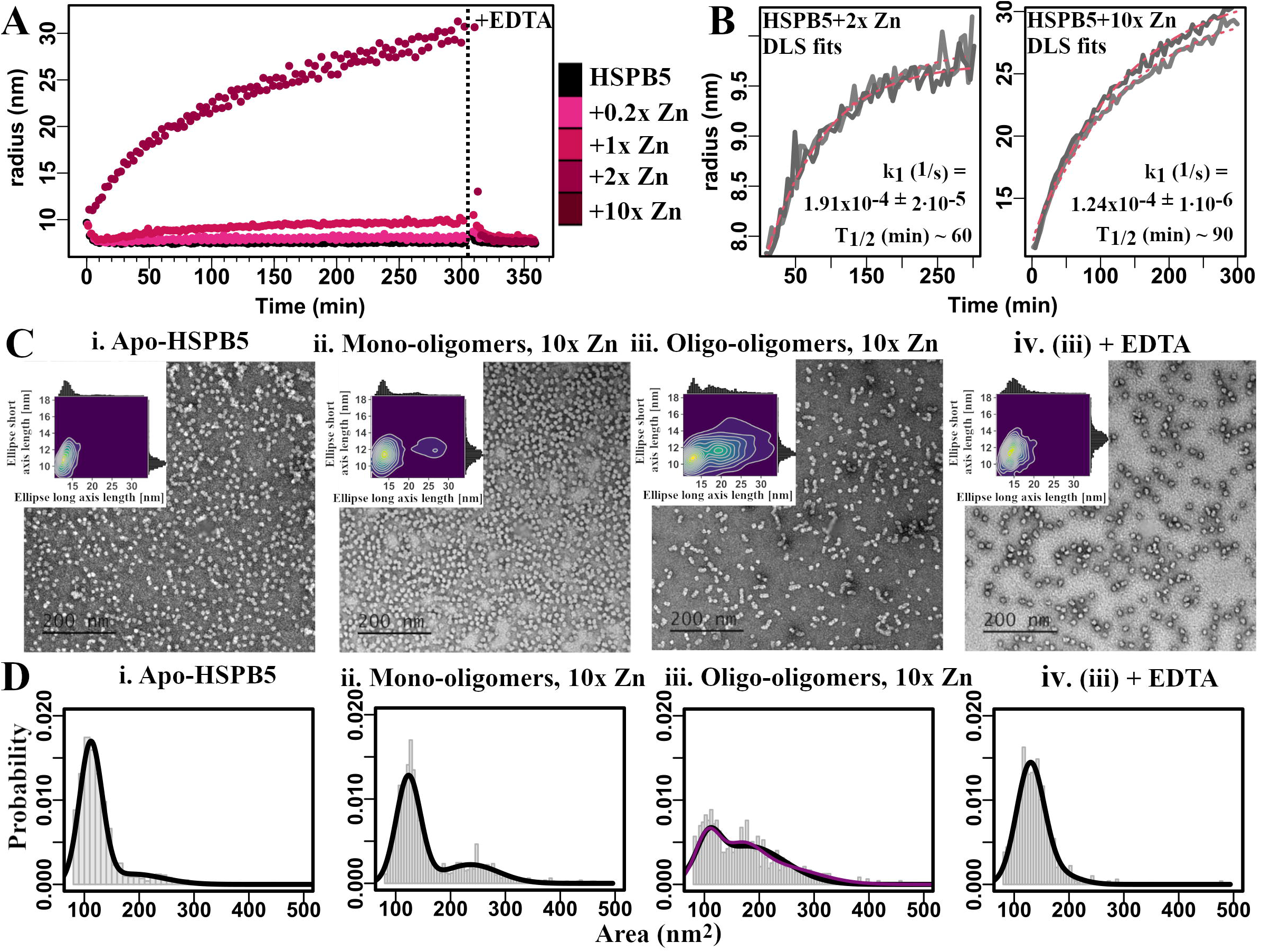
Zinc-induced increase in HSPB5 size is time-dependent and reversible. A. Average hydrodynamic radius as measured by DLS was monitored over time at 37°C following addition of 0.2x, 1x, 2x, 10x zinc (pink color gradient). Reversibility was assessed by addition of EDTA. B. Fits of DLS time courses for HSPB5 following addition of 2x and 10x zinc. Note difference in y-axis. C. Conformational rearrangement of HSPB5 in the presence of zinc visualized by negative-stain EM. i. Apo-HSPB5. ii. Mono-oligomers ((-) incubation), 10x Zn, iii. Oligo-oligomers ((+) incubation), 10x Zn, iv. (iii) + EDTA. Insets: EM images were processed and ellipsoids were fitted into the particles. Plots show 2D density plots of fitted minimum short and long ellipsoid axes. D. Histograms of area distribution (grey bars) with 2-or 3-Gaussian fits (black or purple, respectively).

The relatively slow time course of zinc-induced effects on HSPB5 raised the question of whether the species formed represent an irreversible soluble aggregate. To test for reversibility, the strong zinc chelator EDTA (K D ∼10^-16^ M (36)) was added to samples containing high molecular weight HSPB5 species. The average hydrodynamic radius rapidly returned to its initial value following addition of EDTA; the reversal is complete in ∼5 minutes at 37 °C (Fig. 2A). The results indicate that zinc-induced changes in HSPB5 oligomer size are fully reversible and that return to the apo-like state occurs in a matter of minutes at physiological temperature.

DLS indicated formation of large HSPB5 species in the presence of zinc, but because DLS is biased toward large particles, we examined their morphology and prevalence by negative-stain EM (Fig. 2C). Zinc-free HSPB5 oligomers appeared symmetric and round, consistent with prior reports (Fig. 2C.i) (19). When applied directly to grids, HSPB5 with 10× zinc looked similar (referred to here as “mono-oligomers”) (Fig. 2C.ii). After 3 h at 37 °C, zinc-containing HSPB5 formed clusters of 2–5 oligomers in various arrangements (“oligo-oligomers”) (Fig. 2C.iii). Addition of EDTA to these clusters reverted them to zinc-free HSPB5-like species (Fig. 2C.iv), consistent with the DLS results.

Images were analyzed by quantitative particle analysis (37) using two approaches: ellipsoid fitting and surface area calculation for each particle (Table S2). For zinc-free HSPB5, particles appeared spherical, with short and long axis lengths centered around 11 nm, and >80% classified as mono-oligomers, consistent with the known polydisperse size distribution (Fig. 2C.i). Zinc-containing HSPB5 without incubation showed similar axis lengths, but ∼30% of particles exhibited doubled long axes and surface areas, suggestive of two oligomers in contact, while ∼70% remained mono-oligomers (Fig. 2C.ii). After incubation with zinc, particles displayed broader distributions in both short and long axes, with greater expansion along the long axis, consistent with end-on-end oligomer associations. In these samples, the mono-oligomer population dropped below 30% (Fig. 2C.iii, Table S2). Finally, addition of EDTA to zinc-induced clusters restored particles to symmetric shapes comparable to zinc-free HSPB5 (Fig. 2C.iv).

The EM data are congruent with the kinetic parameters obtained from DLS data (Fig.2B). Taken together they show that HSPB5 undergoes a slow zinc-dependent transition to higher-order oligo-oligomers that dissociate rapidly in the presence of a competitive zinc-binding species. We propose that the oligo-oligomers are composed of two or more oligomers and are stabilized by Zn^2+^ at their inter-oligomeric interfaces.

### Zinc-dependent conformational and dynamic changes in of HSPB5 oligomers

To assess zinc effects on HSPB5 conformation, far-UV circular dichroism (CD) spectra were collected for apo-HSPB5 and oligo-oligomer samples formed with 1×, 2×, or 10× zinc. Compared to apo-HSPB5, oligo-oligomers showed a pronounced increase in negative ellipticity with a shift to lower wavelength, consistent with increased disorder (Fig. 3A). Time-course CD spectra following zinc addition confirmed that these changes occur over the same period as the slow zinc-dependent alterations observed by DLS and EM. Again, the CD changes were fully reversible with EDTA (Fig. S8). Thus, zinc-driven formation of oligo-oligomeric HSPB5 is accompanied by increased secondary structure disorder.

**Figure 3.**
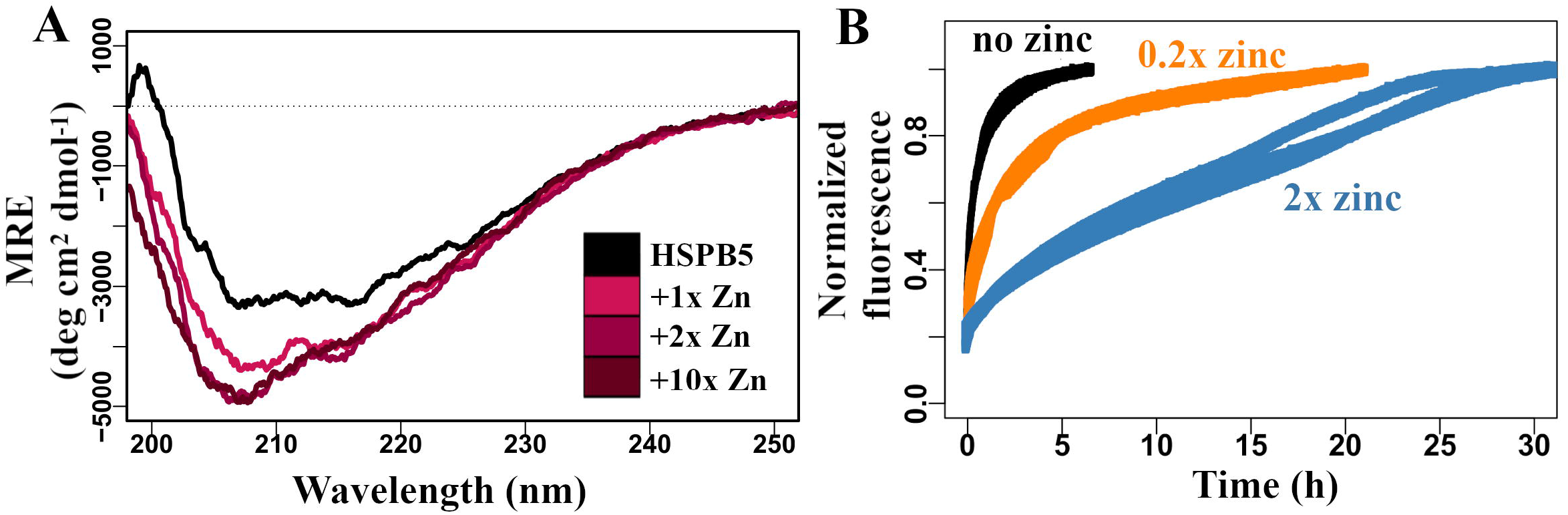
Zinc induces changes in HSPB5 conformation and dynamics. A) Secondary structure as monitored by CD at 25°C. 10 μM HSPB5 was incubated at 37°C for 3h in the absence or presence of zinc (0x (black), 1x (pink), 2x (dark pink), 10x (maroon)); B) Subunit exchange as a function of time at 37°C in the presence of 0, 0.2x, and 2x. Data shown in triplicate.

HSPB5 subunits exchange between oligomers, a property thought to underlie chaperone activity (38), prompting us to ask whether zinc binding affects this process. Subunit exchange kinetics at 37 °C were measured using a fluorescence assay (39) in the absence and presence of Zn^2+^ (Fig. 3B, Table S3). Without zinc, the exchange rate was ∼3 × 10^−4^ s^−1^, corresponding to one subunit per hour. Addition of sub-stoichiometric zinc (0.2 mol equiv per subunit, ∼5 Zn^2+^/oligomer) slowed exchange 6-fold to ∼7 × 10^−5^ s^−1^, or one subunit every 4 hours. Equilibration with 2× zinc (∼48 Zn^2+^/oligomer) produced an even greater 50-fold reduction, yielding a rate of ∼6 × 10^−6^ s^−1^, or one subunit every 50 hours. Thus, zinc binding effectively halts subunit exchange, “freezing” subunits within HSPB5 oligomers.

### Structural basis for zinc binding by HSPB5

HSPB5 and HSPB4 share six structurally analogous histidines, but only HSPB5 displays high-affinity zinc binding and zinc-dependent transitions. What structural features underlie this difference? As a point of reference, we examined zinc-binding proteins with experimentally determined structures that use ≥3 histidines to coordinate zinc. In a representative structure (carbonic anhydrase, PDB IM1 (40), His side chains arrange tetrahedrally around Zn^2+^ with inter-imidazole distances of 3.3–3.6 Å (Fig. S9A). In the highly conserved ACDs of HSPB5 and HSPB4, each dimer contains 10 His positioned on two faces of the domain (Fig. 4A). On one, His101 and His119 face each other on neighboring b-strands with His–His distances of 3.9 Å, but the spacing between pairs in the dimer is >10 Å, precluding coordination of a single zinc ion without major distortion. On the opposite face, His83 and His104 are separated by 5.0 Å (PDB: 2N0K (20)) and could potentially approach one another through sidechain rotation. Thus, the ACD dimers lack pre-formed zinc-binding sites but could act as incomplete templates that could be completed by residues contributed by the highly plastic NTR.

**Figure 4.**
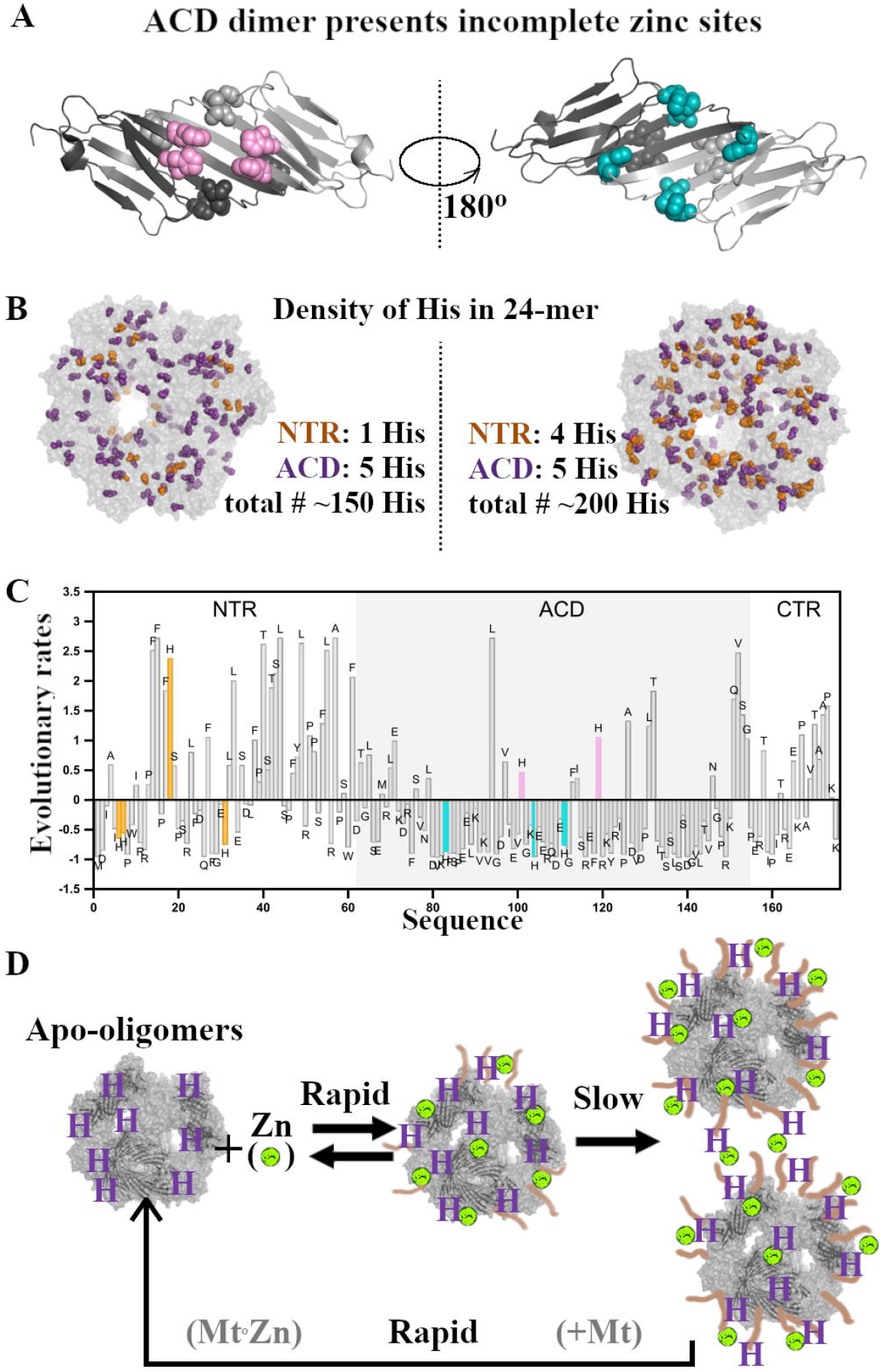
Structural insights into HSPB5 zinc binding. A. HSPB5 ACD dimer (PDB 2N0K) showing histidines: (*Left*) H101 and H119 in pink (*Right*), H83, H104, and H111 in cyan. B. Envelopes illustrating histidine density in a 24-mer. (*Left*), an envelope with 6 His per oligomer (as in HSPB4’s NTR and ACD); (*Right*) an envelope with 9 His per oligomer (as in HSPB5). C. Normalized evolutionary rate profile of human HSPB5. Each bar represents the site-specific evolutionary rate estimated from a multiple sequence alignment of vertebrate orthologs. Rates below zero indicate slower than average evolution, suggesting stronger selective constraints. NTR His are in yellow; ACD His are in pink and cyan, as in Panel A. D. A possible mechanism of HSPB5-mediated zinc buffering during stress. Zinc (lime spheres) can bind to HSPB5 by coordination with His. Prolonged binding leads to conformational rearrangements in HSPB5 and the formation of oligo-oligomers. Zinc can be rapidly released from HSPB5 when canonical zinc-coordinating proteins (e.g., metallothioneins, MTs) are available.

Zinc coordination has been observed in bovine HSPB4 ACD crystals (PDB: 3L1E (41)), where a zinc is coordinated by three His from three separate chains, two are analogous to HSPB5 histidines and one is in the HSPB4-specific CTR (Fig. S9B). In a cryo-EM reconstruction of an HSPB4 oligomer (42), the single NTR His (His7) from one chain is proximal to an ACD His (equivalent to HSPB5 His111) from another. Similarly, in a cryo-EM model of HSPB5 (PDB: 2YGD (43)), multiple clusters of NTR His are observed: (1) His6/His7 of one chain with His31 of another, (2) His6/His7 with His18 of another, and (3) His31 from three different chains cluster together. Although these reconstructions represent a single oligomeric state within a quasi-ordered, polydisperse, and dynamic ensemble, they suggest that oligomer assembly can juxtapose histidines to form zinc-binding clusters (illustrated in Fig. 4B). Functional evidence supports this idea: introducing a single His into HSPB4 at positions equivalent to HSPB5 His6 or His31 confers the biphasic zinc-dependent transitions characteristic of wild-type HSPB5, but introducing His18 does not. These results imply that an NTR containing at least two His is required for high-affinity zinc binding and the unusual zinc-dependent behavior.

To explore whether this property reflects evolutionary pressure, we performed an ortholog-based evolutionary rate analysis of HSPB5 (Fig. 4C, S10). The method compares amino acid substitution rates across a multiple sequence alignment, yielding normalized values where negative scores indicate slower-than-average evolution. As expected, most NTR residues evolve rapidly, consistent with intrinsic disorder. Remarkably, however, three of the four HSPB5 NTR histidines (His6, His7, His31) evolve unusually slowly, a pattern often associated with functional importance in disordered regions (44). Only His18 evolves faster than average. Notably, the functional substitutions above map to the slowly evolving histidines. Although not definitive proof that HSPB5 evolved its histidine content specifically to support zinc binding, these structural, mutational, and evolutionary observations together suggest that NTR histidines provide the basis for HSPB5’s unique zinc-dependent properties.

## Discussion

Although total cellular zinc concentrations reach the hundreds of micromolar range, nearly all intracellular zinc is protein-bound, with free zinc maintained at pM–nM levels (7, 8). Most zinc ligands in proteins are cysteines (15), and with ∼10% of the proteome requiring zinc, the irreversible loss of Cys-based binding sites during oxidative stress could be catastrophic (5). This raises the question of what mechanisms buffer zinc when thiol-based sites are oxidized and zinc is released, a situation known as “zinc distress” (33). Among the small number of Cys-free zinc-binding proteins, HSPB5 stands out as a stress-responsive chaperone with known roles in delaying protein aggregation. We asked whether HSPB5 could also serve as a zinc “chaperone” of sorts.

Our results reveal that HSPB5 oligomers, the predominant species in cells, bind zinc with nanomolar affinity and subsequently undergo slow transitions into larger assemblies we term oligo-oligomers. These appear to represent zinc-bound oligomers stabilized by inter-oligomer cross-bridges that likely involve His residues in disordered NTRs. Importantly, zinc-bound oligomers readily revert to their resting state upon chelation with a higher-affinity ligand such as EDTA. Extrapolating to the cellular context, we propose that high constitutive levels of HSPB5 would allow it to rapidly sequester zinc released under oxidative stress and to gradually transition into zinc-storing assemblies. Because metallothionein synthesis takes 24–48 hours to recover after oxidative inactivation (45), HSPB5 could provide a temporary buffer, lowering free zinc sufficiently to prevent mismetalation and aggregation during this vulnerable period. Although zinc-bound oligo-oligomers may have reduced chaperone activity due to frozen subunits, we estimate that each can bind ∼100 zinc ions, a capacity likely sufficient to protect cells from transient zinc overload (Fig. 4D).

Beyond revealing a potential zinc-storage role, our work highlights the ability of intrinsically disordered regions (IDRs) to coordinate metal ions. In HSPB5, histidines within a disordered region confer high-affinity and high-capacity zinc binding. Importantly, whereas zinc dissociation from most canonical zinc-binding proteins occurs on timescales of days-to-years under non-stress conditions (16, 46), tightly-bound zinc dissociates from HSPB5 in a matter of minutes. Introducing a single additional His into the HSPB4 NTR, which normally contains only one, was sufficient to generate high-affinity binding and zinc-dependent structural rearrangements similar to HSPB5. Remarkably, the context of His matters: substitutions at positions equivalent to H6 or H31 in HSPB5 reproduce the zinc-dependent transitions, whereas substitution at H18 does not (Fig. S5). While the structured ACD also binds zinc, its affinity is much lower. Although the hydrophobic NTR cannot be isolated for direct measurement, results from engineered HSPB4 strongly suggest that zinc binding in HSPB5 arises from the interplay of structured and disordered ligands.

Finally, among the ten human sHSPs, only HSPB5 is His-enriched. Our results suggest that HSPB5’s unusual His content—previously linked to pH-sensitive activation (20, 21) —also underlies its zinc-binding property, highlighting a broader principle in which intrinsically disordered regions provide adaptive, stress-responsive metal-binding functions. These findings imply an unexpected intersection between proteostasis and metal homeostasis and raise the possibility that HSPB5 has evolved to protect cells not only from protein aggregation but also from zinc dyshomeostasis.

### Experimental Procedures (See *Supporting information* for details)

#### Mutagenesis and Protein Preparation

Mutagenesis, expression, and purification of constructs was performed as described previously (47, 48).

#### FluoZin-3 Competition Assay

Competition assays were performed with excess FluoZin-3 (20 μM; Invitrogen) relative to zinc (1.5 μM ZnCl_2_) and increasing concentrations of HSPB5 in Chelex-treated buffer @25°C (34, 35, 49).

#### Isothermal Titration Calorimetry

ITC was performed @25°C using 50 μM protein and 2.5 mM zinc. Analysis was performed with MicroCal PEAQ-ITC software (Malvern Panalytical) and NITPIC (50).

#### Circular Dichroism

CD spectra were collected at 25 °C on 10 μM HSPB5 preincubated with zinc (0x, 1x, 2x, 10x).

#### Dynamic Light Scattering

DLS time-courses were run in duplicate on 10 μM HSPB5 with 0x, 0.2x, 1x, 2x, and 10x zinc (49).

#### Subunit Exchange Assay

Alexa-488 labelled HSPB5 and unlabeled HSPB5 were preequilibrated separately with 0x, 0.2x, 2x zinc. Samples were mixed immediately prior reading on a Horiba Fluorolog-3 (39).

#### Electron Microscopy

Micrographs of 1 μM HSPB5 in the presence of 0x, 1x, 2x, or 10x zinc stained with uranyl formate (51) were collected on a FEI Morgagni 100 kV TEM.

#### Evolutionary Analysis

Vertebrate orthologs of human HSPB4 and HSPB5 were retrieved from the NCBI Orthologs database (52). Multiple sequence alignments were generated with Clustal Omega, implemented in UGENE v52.1 (53). Site-specific evolutionary rates were estimated using Rate4Site (54) with default parameters. (55, 56)

## Abbreviations

sHSP: (small heat shock protein)
FZ3: (FluoZin-3)
NTR: (N-terminal region)
ACD: (alpha- crystallin domain
CTR: (C-terminal region)
ITC: (isothermal titration calorimetry)
DLS: (Dynamic Light Scattering)
CD: (Circular Dichroism)
EDTA: (Ethylenediaminetetraacetic acid)
EM: (electron microscopy)
His: (Histidine)
Cys: (Cysteine)

## Acknowledgements

N. Stone for help with protein purification. L. Tuttle for careful reading of the manuscript. U. Jakob for advice on FZ3 competition assays. D. Wilburn for helpful discussions. T. Hinds for help with ITC experiments.

## Funding and additional information

Funding was provided by the National Eye Institute: 2 R01 EY017370 (R.E.K.) & T32 EY07031 (M.K.J.), National Institute on Aging: T32 AG066574 (M.K.J.). and Universidad Nacional de Quilmes (PUNQ918/22). MSF is a researcher at the National Scientific and Technical Research Council (CONICET), Argentina. The content is solely the responsibility of the authors and does not necessarily represent the official views of the National Institutes of Health.

## Conflict of Interests

The authors declare that they have no conflicts of interest with the contents of this article.

## Notes

### Competing Interest Statement

The authors have declared no competing interest.

