## Supplementary_Info for "Small heat shock protein HSPB5 uses disorder to bind zinc with high affinity"

Materials include:

3 tables

10 figures

Extended Experimental Procedures

### Extended Experimental Procedures

#### Mutagenesis and protein preparation

All single residue mutants of FL-HSPB5 and HSPB4 were prepared by side-directed mutagenesis using PfuTurbo DNA Polymerase (Agilent). Expression and purification of HSPB4, HSPB5, and His->Ala HSPB5 mutants were performed as described previously (1, 2).

To ensure lack of binding between buffer and zinc ions, MOPS buffer was used, as it is established that it does not bind zinc. Buffer conditions used: 25 mM MOPS, 150mM NaCl, in pH 7.5 unless specified otherwise. Zinc was added from 20 mM stocks of zinc chloride salts dissolved in water.

#### Preparation of HSPB5-Zinc oligomers

HSPB5 is highly sensitive to solution conditions. Typically, the response to changes in conditions is very fast (as in the case of pH). In comparison, responses of HSPB5 to zinc is a slower process. We monitored zinc-induced transitions by DLS and SEC and found it necessary to incubate HSPB5 with zinc at physiological temperature (37°C) for ~3h; the transitions are not observed at room temperature or on ice (Fig. S6&S7). We used these insights to prepare distinctly different species of HSPB5 for study. To assess HSPB5 conformation, we prepared two zinc-bound states (mono-oligomers and oligo-oligomers) that correspond to the initial conformation upon zinc-binding and the final conformation. To prepare mono-oligomers state, HSPB5 was incubated for 3h without zinc, followed by addition of zinc and storage on ice until the start of an experiment. To prepare oligo-oligomers, HSPB5 was incubated in the presence of zinc for 3h at 37°C. After incubation, samples were placed on ice until an experiment was performed (maximum few hours). We tested that EDTA can strip zinc from oligo-oligomers by monitoring the decrease in size via DLS and learned that the recovery process is temperature-dependent: at 37°C return to apo-state takes ~5 min and at 25°C recovery takes ~40-50 minutes.

#### FluoZin-3 zinc competition assay

To avoid zinc contamination: buffers (25mM HEPES, 150mM NaCl, pH 7.5) were made with optima water and chelexed overnight. Competition assays included excess FluoZin-3 (20 μM; Invitrogen) relative to zinc (1.5 μM zinc in the form of $ZnCl_2$) to which increasing amounts of HSPB5 (0, 1, 2, 2.5, 5, 10, 25, 50, 75, 100, 150μM) were added. HSPB5 was incubated for 1h @37°C before the experiment without zinc or FluoZin-3. Assays were performed at 25°C to avoid conformational rearrangement of HSPB5 upon zinc binding. Titration points were performed in duplicate. Two curves are shown for each data point. Experiments were performed on a BMG LABTECH CLARIOstar Plus using low volume black 96 well plate (Corning). Data were fitted assuming the following processes occurring in the sample:

$$Zn + FZ3 \rightarrow Zn - FZ3$$

$$Zn + HSPB5 \rightarrow Zn - HSPB5$$

And solved using quadratic equation for the Kd of sHSPs similarly to Jakob *et al.* (3)

Where:

$$K_{D\ (FZ3)} = \frac{[FZ3]_{free} * [Zn]_{free}}{[FZ3]_{bound}}$$

$$K_{D\ (HSPB5)} = \frac{[HSPB5]_{free} * [Zn]_{free}}{[HSPB5]_{bound}}$$

And assuming:

$$[FZ3]_{free} = [FZ3]_{total} - [FZ3]_{bound}$$

$$[HSPB5]_{free} = [HSPB5]_{total} - [HSPB5]_{bound}$$

$$[HSPB5]_{bound} = [Zn]_{total} - [HSPB5]_{bound}$$

The resulting quadratic equation had a following form:

$$[FZ3]_{bound} \sim \frac{-b + \sqrt{(b^2 - 4ac)}}{2a}$$

$$a = K_{D\ (FZ3)} - K_{D\ (HSPB5)}$$

$$b = K_{D\ (FZ3)} * [B]_{tot} - K_{D\ (HSPB5)} * [Zn]_{total} + K_{D\ (HSPB5)} * ([FZ3]_{total} - [Zn]_{total})$$

$$c = [FZ3]_{total} * K_{D\ (HSPB5)} * [Zn]_{total}$$

The data were fitted using Levenberf-Marquardt Nonlinear Least-Squares Algorithm with minpack.lm R package (4), using a value of 9.1nM for $K_{d(FZ3)}$ (5).

Isothermal calorimetry

ITC measurements were performed on a Microcal VP-ITC calorimeter (Malvern Panalytical) at 25°C. Proteins in the ITC cell were at 50 µM in 50 mM MOPS, 150mM NaCl. Zinc chloride (2.5 mM) was used as titrant. Binding curves at pH 7.5 and 6.5 were highly similar, so experiments were performed at pH 6.5 (Fig. S1) to avoid solubility issues. Injection volume was 3uL and injections were in 4-min intervals. Data were analyzed using Microcal PEAQ-ITC analysis software (Malvern Panalytical). Baselines for the plots were calculated using NITPIC software (6).

Circular Dichroism

Circular dichroism spectra were collected on a Jasco J-1500 CD spectrometer with a 1 nm bandwidth on proteins dissolved in 1 mM MOPS, 6 mM NaCl at pH 7.5. For time-course experiments, 2.5 µM HSPB5 was preincubated at 37°C for 1h before 30x zinc addition. After recording the apo-HSPB5 spectrum 75 µM zinc was added and wavelength scans were performed at multiple timepoints for 3h, after which 0.5 mM EDTA was added and another spectrum was recorded. For monitoring zinc dose-dependent changes in the HSPB5 spectrum 10 µM protein was preincubated for 3h at 37°C in the presence or absence of zinc (0x, 1x, 2x, 10x), spectra were recorded at 25°C. All the data shown have a high-tension below 600 V.

Dynamic Light Scattering

Dynamic Light Scattering was performed on a DynaPro Plate Reader III from Wyatt Technology Corporation using Aurora 384-well plate sealed with film to prevent evaporation. Data were analyzed with DYNAMICS 7.8.2 software provided with the instrument. Experiments to monitor the time-course of protein size were performed in duplicate using 10 μM HSPB5 variants in 25 mM MOPS, 150 mM NaCl, pH 7.5 with 0x, 0.2x, 1x, 2x, and 10x zinc addition. Proteins were filtered before experiments using 0.22 μm filters. All samples were incubated for 1h at 37°C before being added to a preheated plate. DLS measurements were started immediately and were performed at 37°C or 25°C (control). EDTA (1 mM) was added to all wells after samples reached a plateau. The increase in size was later analyzed by assuming first order reaction kinetics and fitting to $y = C1 – C2*exp(-k_1t)$ using minpack.lm R package (4). C1 represents the maximum radius and $k_1$ is the apparent rate constant.

Subunit Exchange measured by fluorescence

Alexa Fluor 488 maleimide labeling protocol was described previously (7). 15 μL of 2 μM of S153C HSPB5 labeled with Alexa (81% label efficiency) and 10 μM wild type, unlabeled HSPB5 (135 μL) were preincubated at 37°C for 3h with or without zinc (0, 0.2x, 2x zinc) before starting an experiment. The samples were mixed in a 37°C preheated cuvette just prior to starting an experiment. Experiments were performed on a Horiba Fluorolog-3. Excitation and emission wavelengths for Alexa Fluor 488 maleimide 498/518 nm were used. The increase in size was later analyzed by assuming first order reaction kinetics and fitting to $y = C1 – C2*exp(-k_1t)$ (4). All the experiments were performed in triplicate.

Electron microscopy

Negative-stain TEM was performed using the single droplet procedure at ambient temperature (8). Micrographs were collected on FEI Morgagni 100 kV TEM equipped with Gatan Orius Camera. 20 μM HSPB5 was incubated with 0 μM, 20 μM, 100 μM, or 200 μM zinc at 37°C for 3h. Additional samples of 20 μM HSPB5 incubated with 200 μM zinc on ice for 3h, with zinc added after incubation at 37°C and 20 μM HSPB5 incubated with 200 μM zinc for 3h then stripped with 1mM EDTA and incubated at 37°C for 30 minutes were prepared. Buffer conditions in 25 mM MOPS, 100 mM NaCl, pH 7.5. Samples were diluted 20x for grid preparation and stained with uranyl formate.

Images were analyzed by quantitative particle analysis using Fiji (9) in two ways: 1) particles were fit to an ellipsoid and 2) surface area was calculated for each particle (Table S2). The analysis was performed on at least 500 particles from each grid. We compared particle width vs length using the short and long ellipse axis lengths from the ellipsoid fits (insets in Fig. 2C). The surface area analysis of particles was used to estimate relative populations of species under each condition. Histograms of particle surface areas were fitted to a two-Gaussian model (Fig. 2D and Table S2); the oligo-oligomer data were also fit to a three-Gaussian model (Table S2). The area under each curve represents the relative population.

Size exclusion chromatography (SEC)

SEC experiments on a Superose 6 10/300 GL column (GE Life Sciences) were performed on GE AKTA Purifier. Injection volume was 100μM. The zinc concentration effect was accessed by adding different amounts of zinc (0, 1x, 2x) to HSPB5 and incubating for 3h at 37°C. For assessing the importance of the incubation effect on size changes of HSPB5 in presence of zinc 100 μM HSPB5 with 1x zinc was used in 150 mM NaCl, pH 7.5. Different length of incubation of HSPB5 with zinc @ 37°C: 0, 1h, 3h incubation.

Evolutionary analysis

To obtain the site-specific evolutionary rate profiles of human HSPB5 and human HSPB4, vertebrate orthologous sequences were retrieved from the NCBI Orthologs database (https://www.ncbi.nlm.nih.gov/gene), using one representative protein per species. The queries were based on the human genes CRYAB (Gene ID: 1410) and CRYAA (Gene ID: 1409), which encode HSPB5 and HSPB4, respectively. Only orthologous sequences were included to ensure that the dataset reflected the evolutionary history of a single ancestral gene, allowing meaningful estimation of relative evolutionary constraints at each site along the protein sequence. Manual curation was performed to retain only full-length sequences and to exclude those containing undefined residues (e.g., “X”). The final datasets included 369 sequences for HSPB5 and 424 for HSPB4.

Multiple sequence alignments were generated using Clustal Omega, as implemented in the UGENE software (v52.1) (10). Rate4Site program (11) was used with default parameters (JTT as amino acid substitution matrix and neighbor-joining method to reconstruct the phylogenetic tree) (12, 13) to estimate normalized evolutionary rates at each aligned position. Human sequences were used as reference for both analyses (HSPB5: UniProt P02511, HSPB4: UniProt P02489).

**Comments about HSPB5 and evolutionary rate estimation:**

Residues that evolve at slower rates are typically under positive selective pressure, often reflecting either structural or functional constraints. Given the intrinsically disordered nature of the NTR, the observation that histidine residues within this region exhibit evolutionary rates lower than the sequence average suggests a potential functional relevance.

In contrast, histidines within the ACD are part of a domain that is highly conserved among sHSPs, and residues in the β-strand regions are generally expected to evolve slowly due to their involvement in maintaining domain structure through stabilizing interactions. Histidines located in the ACD have a range of pKa values, suggesting their involvement in different functional aspects (14).

His101 and His119 exhibit scores higher than the sequence average. Both face the exterior surface of the ACD dimer in the 2N0K structure and their imidazole rings are close to each other. Notably, in the MSA, His119 is conserved in only 208 out of 369 sequences (56.4%), whereas His101 is conserved in 345 out of 369 (93.5%). The most frequent substitutions for His119 are asparagine, serine, and glutamine. This suggests that, although in human HSPB5 these two histidines are spatially close and interact within an ACD, their proximity is not strongly conserved across vertebrates.

In contrast, His104, which points toward the inner surface of the ACD, and His83 and His111, located in loop regions, appear to participate in a broader network of interactions via their side chains. His104 is 100% conserved across the MSA and aligns with His100 in HSPB4, which directly coordinates a $Zn^{2+}$ ion in the 3L1G structure. Furthermore, the functional relevance of HIS104 in mediating interface stability and regulating oligomer structure in response to acidosis has been demonstrated (14), suggesting its evolutionary conservation reflects a critical functional role that should be considered when interpreting its score.

Finally, His7, the only His residue within the HSPB5 NTR that is also present in HSPB4, is conserved in 71.3% of the sequences and is replaced by Asn in 27.1%. Notably, while HIS7 is highly conserved in HSPB4 (96.7%), this paralog does not exhibit the high-affinity zinc binding observed in HSPB5. High-affinity zinc binding and zinc-induced transitions are only recovered in HSPB4 when mutations equivalent to HSPB5’s His6 and His31 are introduced.

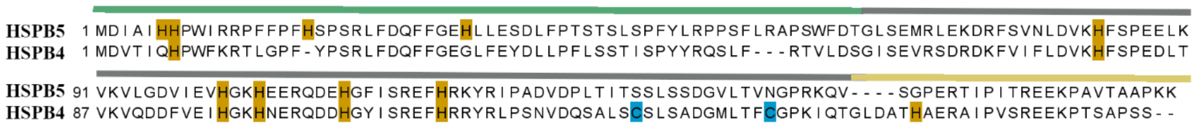


Figure S1 Sequence alignment of HSPB5 and HSPB4. Boundaries of the domains of HSPB4 and HSPB5 are highlighted (green-NTR, grey -ACD, yellow-CTR) above the sequence. Histidine residues are highlighted in orange. Cysteines in HSPB4 are shown in blue. For the ITC experiments the cysteines were substituted with serine.

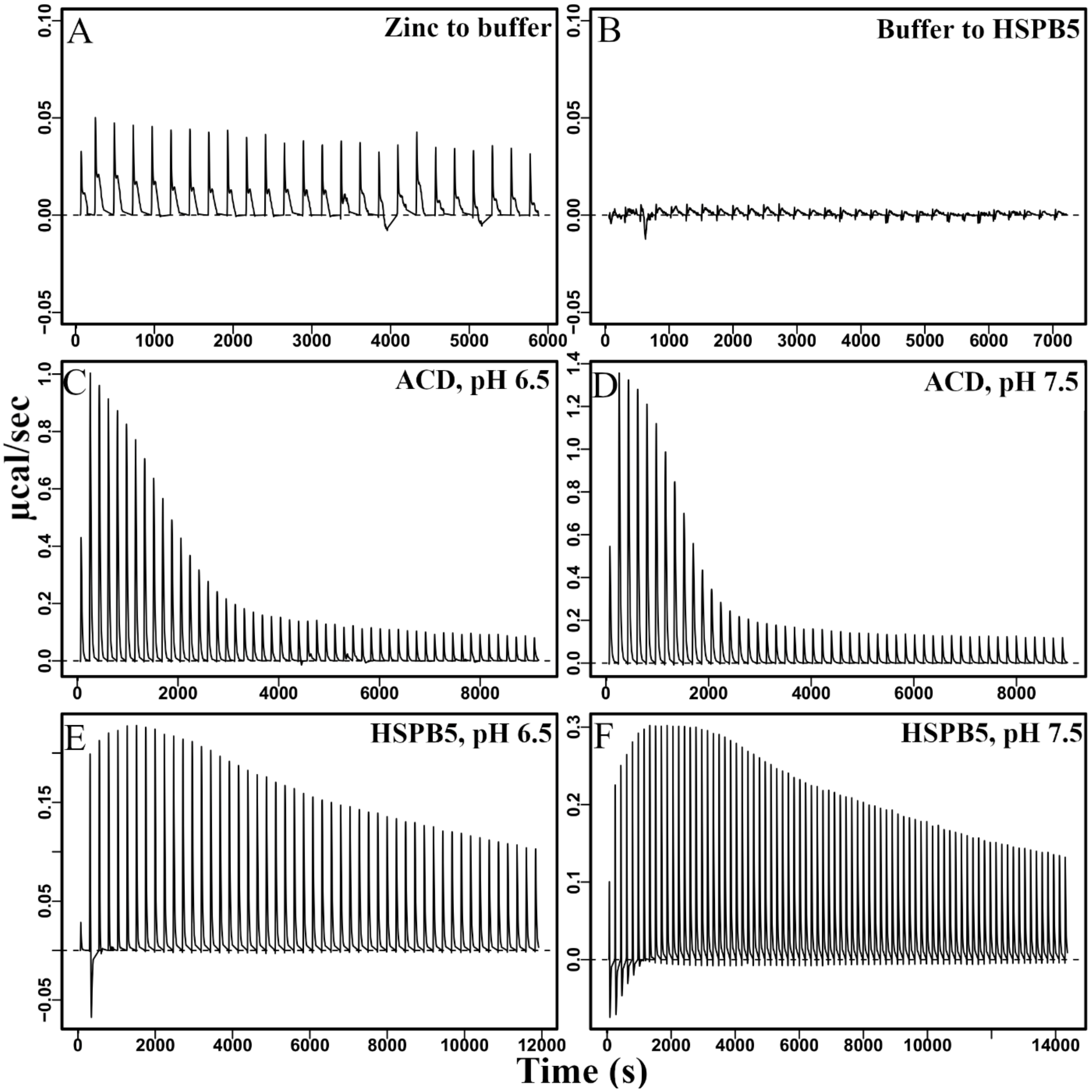


Figure S2 Raw ITC profiles. Controls and WT constructs in pH 7.5 and 6.5

| | Protein | $\Delta H_1$ (kJ/mol) | $\Delta G_1$ (kJ/mol) | $-T\Delta S_1$ (kJ/mol) | $\Delta H_2$ (kJ/mol) | $\Delta G_2$ (kJ/mol) | $-T\Delta S_2$ (kJ/mol) | Red. Chi-Sqr. $(kJ/mol)^2$ |
|---|---|---|---|---|---|---|---|---|
| 1 | HSPB5 pH 7.5 | -173 | -26.6 | 146 | 6.25 | -20.5 | -26.8 | $6.20 \times 10^{-3}$ |
| 2 | HSPB5 pH 6.5 | -261 | -26.4 | 235 | 5.01 | -19.8 | -24.8 | $9.90 \times 10^{-3}$ |
| 3 | HSPB4 | 0 | 0 | 0 | 5.1 | -23.6 | -28.7 | $5.50 \times 10^{-3}$ |
| 4 | HSPB4-3His | -3.8 | -26.5 | -22.7 | 4.92 | -19.5 | -24.5 | $5.40 \times 10^{-3}$ |
| 5 | ACD, pH 6.5 | 0 | 0 | 0 | 17.7 | -29.3 | -46.9 | $6.10 \times 10^{-2}$ |

Table S1 Fit parameters for ITC profiles.

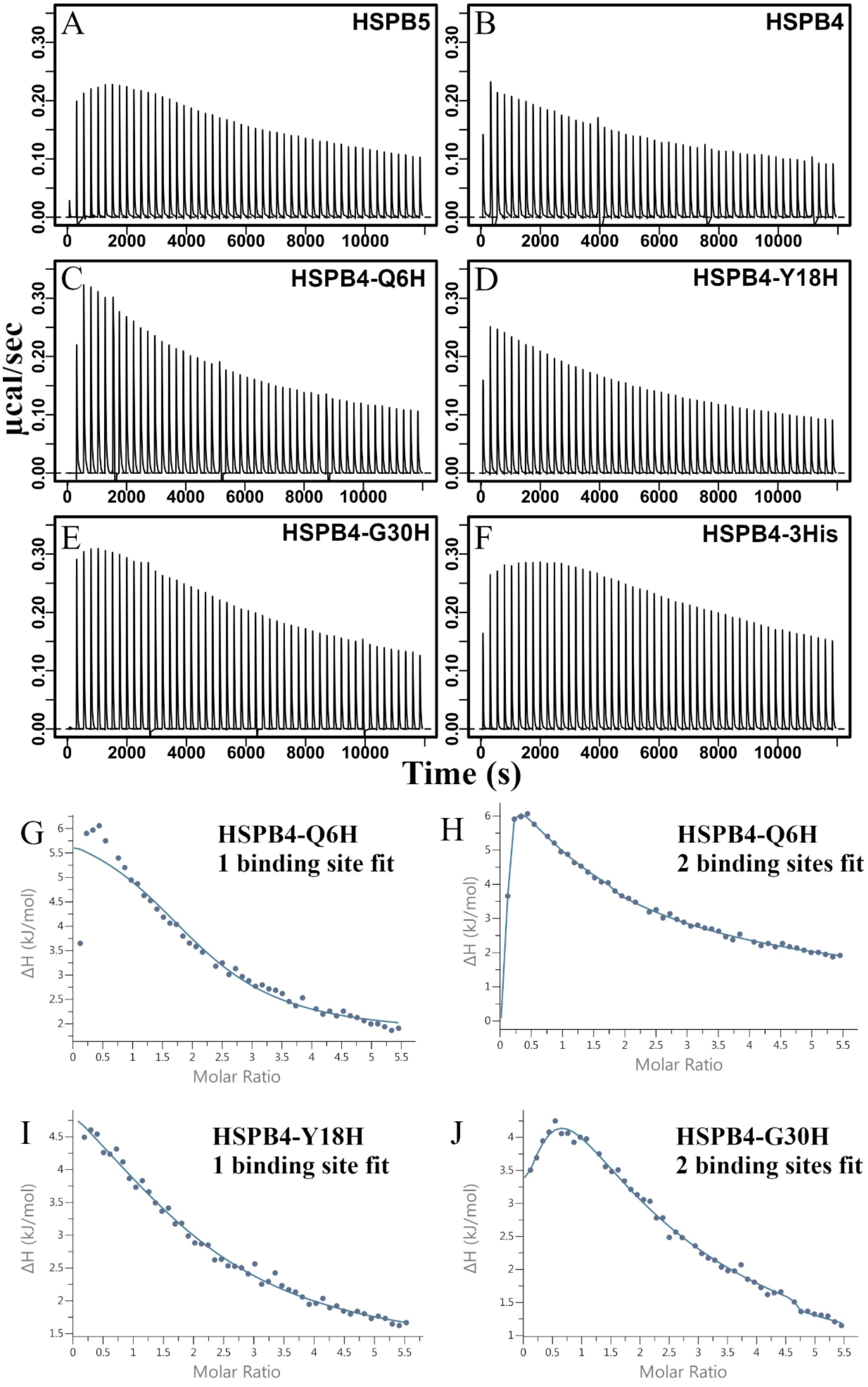
A HSPB5
B HSPB4
C HSPB4-Q6H
D HSPB4-Y18H
E HSPB4-G30H
F HSPB4-3His
µcal/sec
Time (s)
G HSPB4-Q6H 1 binding site fit
H HSPB4-Q6H 2 binding sites fit
I HSPB4-Y18H 1 binding site fit
J HSPB4-G30H 2 binding sites fit
ΔH (kJ/mol)
Molar Ratio

Figure S3 A-F Raw ITC profiles. A. HSPB5, B. HSPB4, C.  HSPB4-Q6H, D. HSPB4-Y18H, E. HSPB4-G30H, and F. HSPB4-3His in which all 3 NTR histidine installed. G-J Isotherms with fitted curves. G. HSPB4-Q6H, single transition fit does not recapitulate the data well. C.  HSPB4-Q6H 2 transition fit, D. HSPB4-Y18H single transition, E. HSPB4-G30H 2 transitions fit.

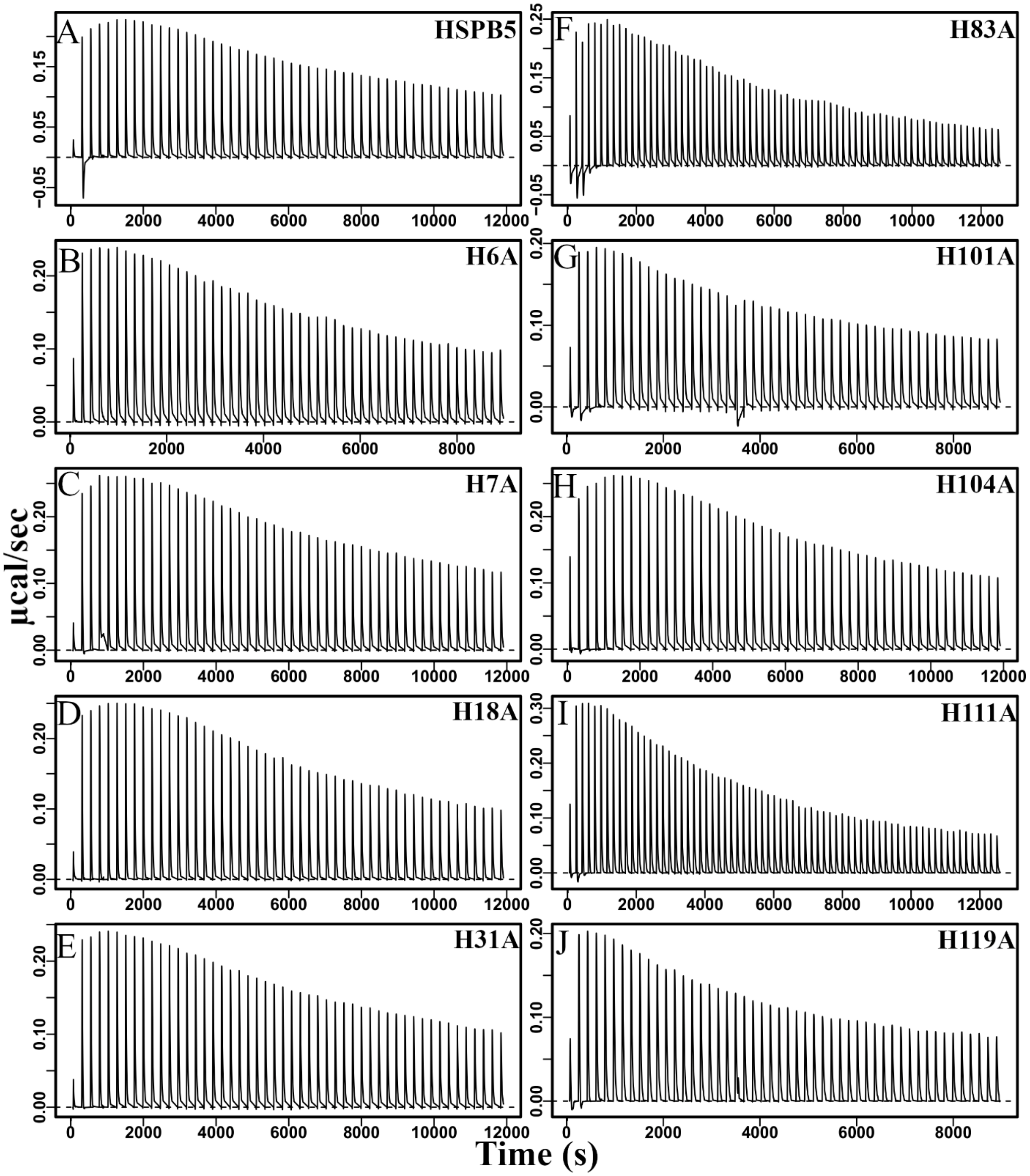


Figure S4 Raw ITC profiles. Histidine-to-Alanine mutants of HSPB5.

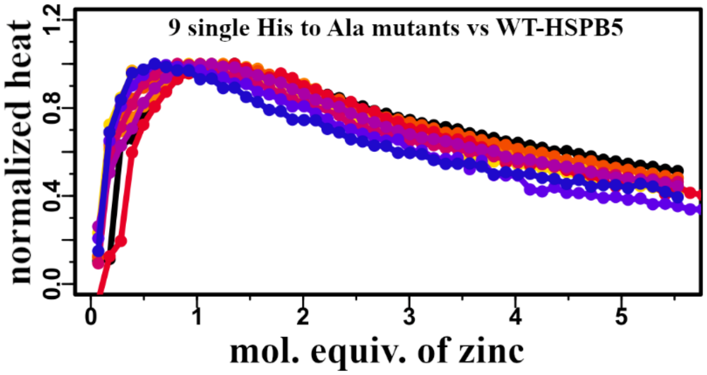


Figure S5 Normalized heat derived from the ITC experiment for His-to-Ala mutants of HSPB5. The overlay of the 9 single His to Ala mutants with WT HSPB5 shows that the biphasic pattern of the HSPB5 isotherm upon zinc binding is retained for every variant.

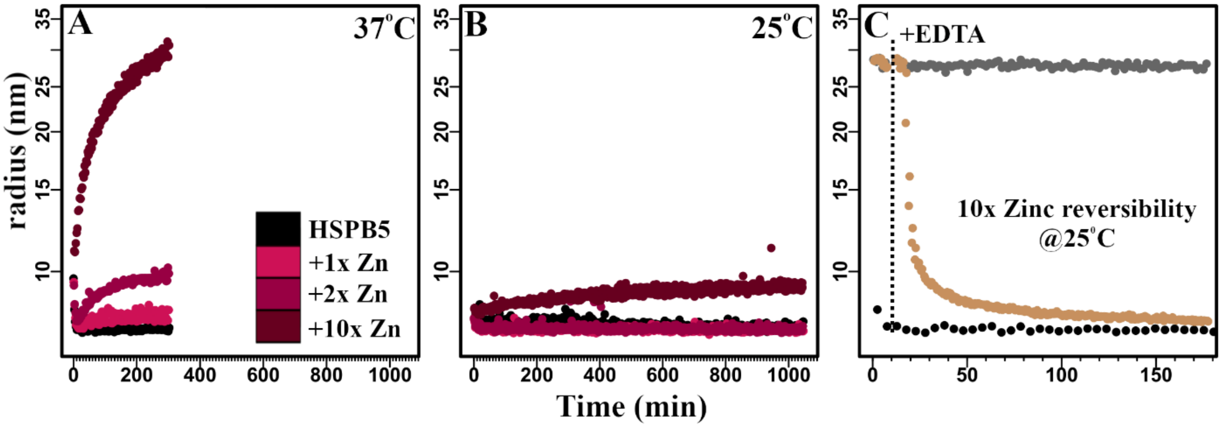


Figure S6 Effect of temperature on the zinc-induced change in HSPB5 size as measured by DLS. A. Change in size of HSPB5 at 37°C (no zinc (black), 1x zinc (magenta), 2x zinc (dark pink), 10x zinc (maroon)) (data from the main text figure 2). B. Change in size of HSPB5 at 25°C (no zinc (black), 1x zinc (magenta), 2x zinc (dark pink), 10x zinc (maroon)). Axes for Panels A and B are shown in the same range to allow direct comparison. The Y-axis is logarithmic to focus on smaller particles. C. The reversibility of formation of high-order species HSPB5 by addition of EDTA at 25°C. Oligo-oligomeric HSPB5 was generated in the presence of 10x zinc at 37°C. Samples were then placed in a plate reader at 25°C and the DLS signal was monitored at 25°C. Complete return to original size required >100 min. Apo-HSPB5 (black), preincubated HSPB5 with 10x zinc (oligo-oligomers) (grey) , preincubated HSPB5 with 10x zinc +EDTA added at the timepoint indicated by dotted line (beige). Experiments performed in duplicate.

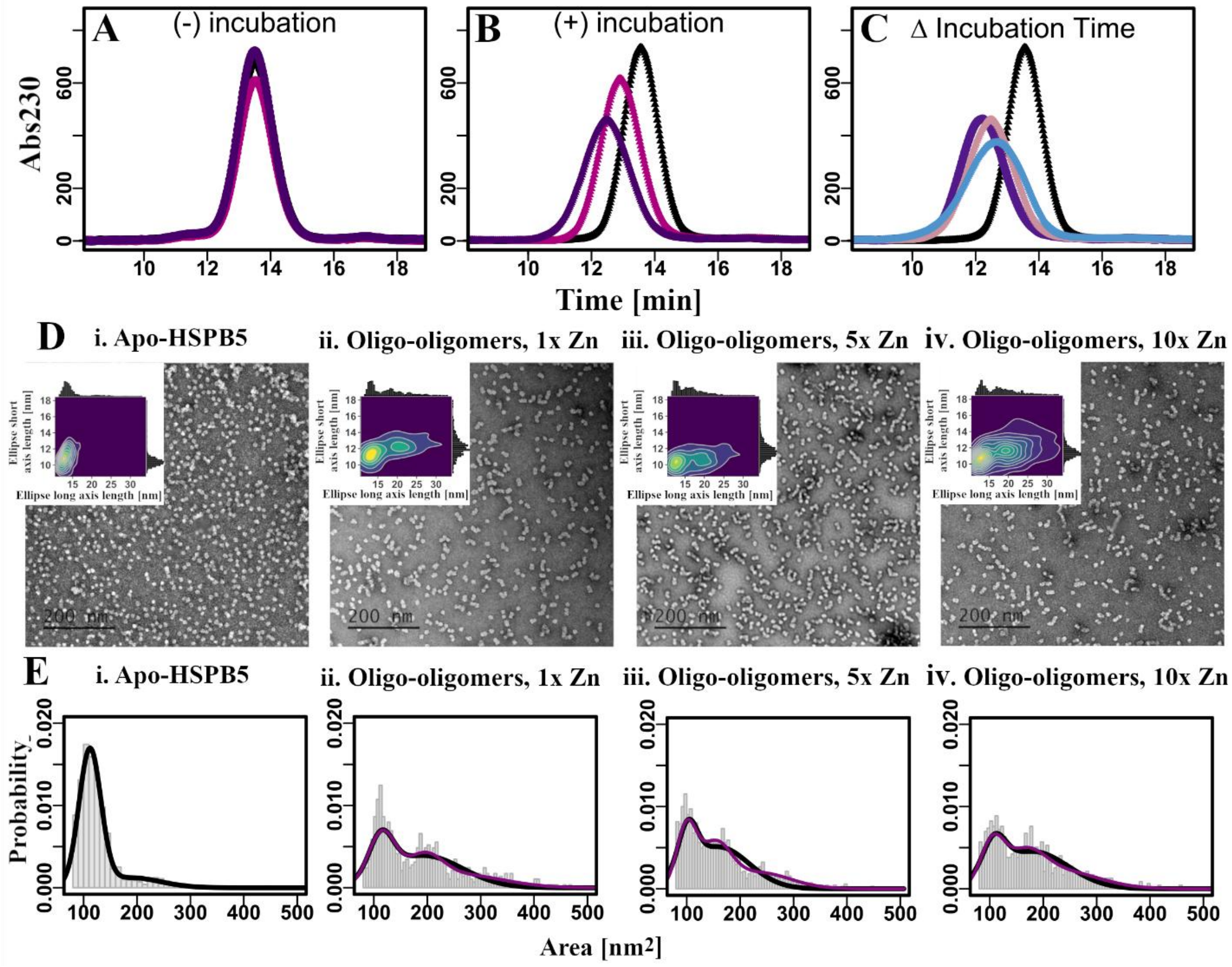


Figure S7 Conformational rearrangement of HSPB5 upon zinc binding. A. SEC performed at room temperature of HSPB5 with 0x (black), 1x (pink), or 2x (dark pink) of zinc on ice, without incubation at 37°C. B. HSPB5 with 0x (black), 1x (pink), or 2x (dark pink) zinc after 3h incubation at 37°C. C. SEC of HSPB5 with 2x zinc after different incubation time @37: 0h (black), 1h (blue), 3h (pink), 18h (purple). D. i-iv. TEM images of HSPB5 in the presence of zinc. i. Apo-HSPB5, ii. Oligo-oligomers, 1x zinc, iii. Oligo-oligomers, 5x Zn., iv. Oligo-oligomers, 10x Zn. Insets: EM images were processed using Fiji and ellipsoid was fitted into the particles. Plots show 2D density plots of fitted minimal short and long ellipsoid axis. Upon incubation with zinc there is an increase in size on long axis of the ellipsoid supporting formation of cross-oligomers interactions. E. Histograms of HSPB5 area distribution (grey bars) with fits to 2 (black line) or 3 (purple line) Gaussians, i. Apo-. ii. Oligo-oligomers, 1x zinc, iii. Oligo-oligomers, 5x Zn., iv. Oligo-oligomers, 10x Zn.

A

| Species | # particles |
|---|---|
| Apo | 998 |
| Mono-oligomers, 10x Zinc | 1636 |
| Oligo-oligomers; 10xZinc + EDTA | 996 |
| Oligo-oligomers, 1x Zinc | 578 |
| Oligo-oligomers, 5x Zinc | 1422 |
| Oligo-oligomers, 10x Zinc | 949 |

B

| Species | % monomers (% AUC 1st Gauss) | % oligomers (AUC 2nd Gauss) |
|---|---|---|
| Apo | 83.3 | 16.7 |
| Mono-oligomers, 10x Zinc | 71.2 | 28.8 |
| Oligo-oligomers; 10xZinc + EDTA | 87.0 | 13.0 |
| Oligo-oligomers, 1x Zinc | 27.5 | 72.5 |
| Oligo-oligomers, 5x Zinc | 24.0 | 76.0 |
| Oligo-oligomers, 10x Zinc | 22.0 | 78.0 |

C

| Species | % monomers (% AUC 1st Gauss) | % dimers (AUC 2nd Gauss) | % oligomers (AUC 3nd Gauss) |
|---|---|---|---|
| Oligo-oligomers, 1x Zinc | 41.5 | 26.4 | 32.1 |
| Oligo-oligomers, 5x Zinc | 33.6 | 36.8 | 29.6 |
| Oligo-oligomers, 10x Zinc | 29.2 | 32.2 | 38.6 |

Table S2 Summary of analysis of EM images. A. Number of particles used for analysis. B. Area under the curve for different species of HSPB5 fitted with 2 gaussians. C. Area under the curve for oligo-oligomers samples fitted with 3 Gaussians.

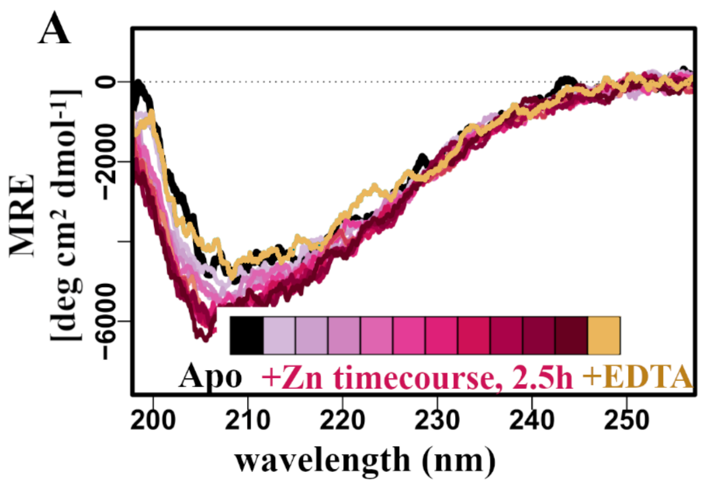


Figure S8 Conformational rearrangement of HSPB5 in the presence of zinc. A) CD spectra of 2.5 µM HSPB5 in the presence of 30x zinc collected in 15 min interval over 2.5h time period at 37°C. Spectrum of HSPB5 in the absence of zinc is in black, time course illustrating conformational rearrangement post-zinc addition is in magenta color gradient. Spectrum collected after addition of EDTA is shown in tan demonstrates reversibility, as it overlays with the black spectrum.

| **Species** | **$k_{ex}$ (1/s)** | | | **Average $k_{ex}$ (1/s)** | |
|---|---|---|---|---|---|
| | #1 | #2 | #3 | | |
| No zinc | $3.5x10^{-4}$ | $3.0x10^{-4}$ | $2.4x10^{-4}$ | $2.9\pm(0.55)x10^{-4}$ | 1 subunit: 1 hour |
| 0.2x zinc | $7.0x10^{-5}$ | $6.7x10^{-5}$ | $7.4x10^{-5}$ | $7.0\pm(0.35)x10^{-5}$ | 1 subunit: 4h |
| 2x zinc | $8.8x10^{-6}$ | $5.5x0^{-6}$ | $3.3x10^{-6}$ | $5.9\pm(2.7)x10^{-6}$ | 1 subunit: 50h |

Table S3 Subunit exchange fit parameters.

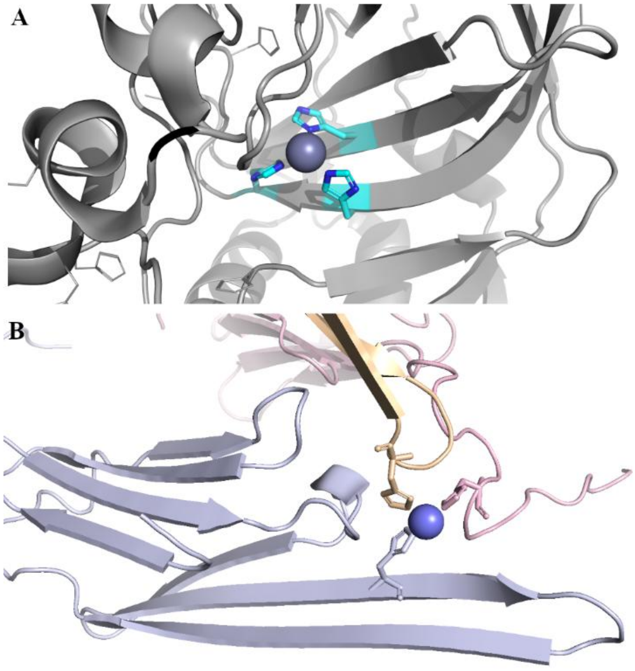


Figure S9 PDB structures of zinc-binding proteins coordinated by three histidine residues. A. α-Carbonic anhydrase from Persephonella marina EX-H1 (PDB: 6IM1). B. ACD of HSPB4, which completes the coordinating sphere using histidines from neighboring crystallographic subunits. The subunits do not form the canonical ACD dimer but are positioned in neighboring subunits, where they interact through the loops. In both structures, zinc (dark blue sphere) is coordinated by histidine residues shown as sticks. For clarity, the three subunits of the ACD are shown in different colors.

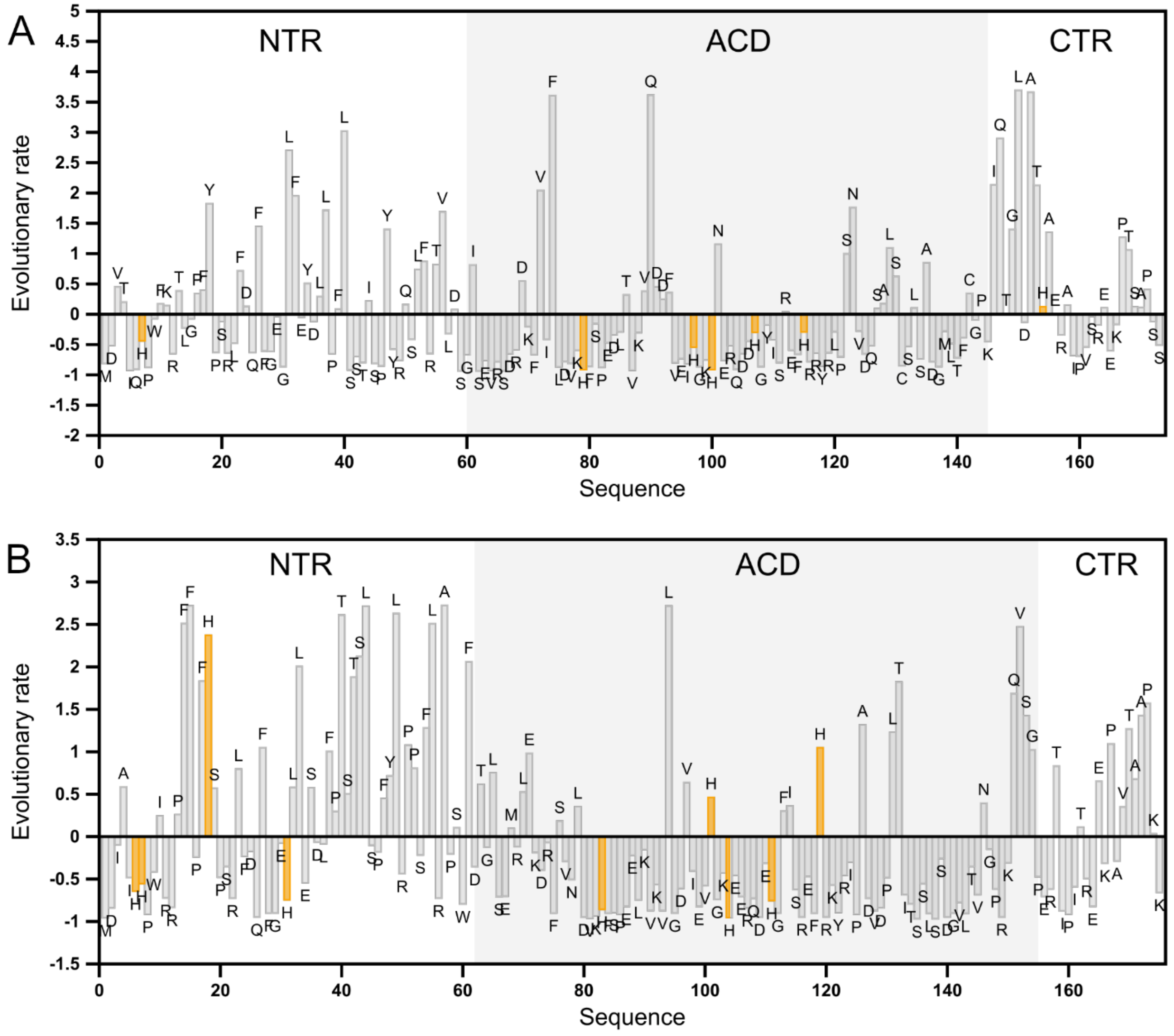


Figure S10 Normalized evolutionary rate profile of human A. HSPB4 B. HSPB5. Each bar represents the site-specific evolutionary rate estimated from a multiple sequence alignment of vertebrate orthologs. Histidine residues are shown in orange. The ACD boundaries are shaded in light grey.
